# Mutation-induced Changes in the Receptor-binding Interface of the SARS-CoV-2 Delta Variant B.1.617.2 and Implications for Immune Evasion

**DOI:** 10.1101/2021.07.17.452576

**Authors:** Prabin Baral, Nisha Bhattarai, Md Lokman Hossen, Vitalii Stebliankin, Bernard S. Gerstman, Giri Narasimhan, Prem P. Chapagain

## Abstract

While the vaccination efforts against SARS-CoV-2 infections are ongoing worldwide, new genetic variants of the virus are emerging and spreading. Following the initial surges of the Alpha (B.1.1.7) and the Beta (B.1.351) variants, a more infectious Delta variant (B.1.617.2) is now surging, further deepening the health crises caused by the pandemic. The sharp rise in cases attributed to the Delta variant has made it especially disturbing and is a variant of concern. Fortunately, current vaccines offer protection against known variants of concern, including the Delta variant. However, the Delta variant has exhibited some ability to dodge the immune system as it is found that neutralizing antibodies from prior infections or vaccines are less receptive to binding with the Delta spike protein. Here, we investigated the structural changes caused by the mutations in the Delta variant’s receptor-binding interface and explored the effects on binding with the ACE2 receptor as well as with neutralizing antibodies. We find that the receptor-binding β-loop-β motif adopts an altered but stable conformation causing separation in some of the antibody binding epitopes. Our study shows reduced binding of neutralizing antibodies and provides a possible mechanism for the immune evasion exhibited by the Delta variant.

## 1. Introduction

The pandemic caused by SARS-CoV-2 infections has surpassed more than 188 million cases worldwide with nearly 4 million fatalities as of July 15, 2021. Several vaccines against the SARS-CoV-2 infections have been developed^1^, with other vaccine candidates in clinical trials^2^. The vaccines authorized for emergency use have been highly effective in reducing the number of cases and deaths^1, 3^. In addition, therapeutic measures are also being pursued in parallel. These include identification of therapeutic small molecules, convalescent plasma, decoy receptors, and neutralizing antibodies^4–13^. Several studies have considered neutralizing antibodies (Abs) that can bind to the virus’ receptor binding domain (RBD)^14–19^ and have advantageous pharmacokinetics and the ability to be produced on a large scale^20^. The primary target for the vaccines or Abs is the receptor binding domain (RBD) of the spike protein, which is responsible for binding of the virus to the human ACE2 receptor on the host cell and facilitating viral entry into the human cells^13, 21, 22^.

While vaccination efforts are ongoing worldwide, new genetic variants are emerging and spreading. Notable variants of concern include the Alpha variant B.1.1.7 (originating in the UK), Beta variant B.1.351 (originating in South Africa), and the Delta variant B.1.617.2 (originating in India). The deadly forms of B.1.617 variants are considered to be largely responsible for the second wave of infections in India, causing a grave health crises with more than 30 millions infections and 400,000 deaths in India alone^23^. The B.1.617.2 Delta variant is considered to be the most infectious of all variants and as of June 2021 has become one of the most transmissible variants with the highest number of reported cases, followed by the B.1.617.1^24, 25^. The Delta variant is fast becoming the most dominant variant in many countries^26^. Fortunately, the current vaccines appear to be effective against many variants of concern, including the Delta variant^27, 28^. However, lack of vaccination coverage worldwide has allowed the virus to spread and continue to evolve, decreasing the chances of quickly ending the pandemic.

Mutations in the RBD can change the ability of the virus spike protein to bind to and enter the host cell. The spike protein of B.1.617.1 has mutations T95I, G142D, E154K, L452R, E484Q, D614G, P681R, Q1071H, and B.1.617.2 has mutations T19R, G142D, Δ156-157, R158G, L452R, T478K, D614G, P681R, and D950N^29^. Since the receptor-binding domain (RBD) of the spike protein is primarily involved in the interaction with ACE2 on the host cell, the RBD mutations L452R/E484Q in B.1.617.1 and L452R/T478K in B.1.617.2 are assumed to play significant roles in infectivity and transmissibility of the virus. Recently, the RBD mutation K417N that is present in the Beta variant has also been identified in the Delta variant, giving yet another novel variant known as the Delta plus variant.

The significant and rapid rise in the number of cases due to infections by the Delta variant in areas that appeared to overcome the earlier surges (e.g. India) or even in areas with high coverage of effective vaccines (e.g. Israel) indicates that the antibodies elicited from vaccines or prior infections by other strains of SARS-CoV-2, or other pathogens through molecular mimicry are not as effective for the Delta variant. Indeed, recent studies^30, 31^ have shown that compared to the previous variants, the Delta variant is not only able to evade immunity conferred by previous infections but is also less sensitive to neutralizing Abs from recovered patients. Some anti-RBD Abs have been shown to have reduced RBD binding, resulting in a 4-fold decrease in the potency of the sera from prior infections and a 3-5 fold decrease in vaccine-generated Abs against the Delta variant^31^. Similarly, it has been shown recently that the mutations in the B.1.427/B.1.429 variants cause reduced or complete loss of sensitivity to RBD-binding antibodies (Abs)^32^. This suggests that, in addition to possibly a high affinity of the RBD to ACE2 or its ability to present itself in an up conformation in the spike trimer^33^, antibody evasion due to RBD mutations may also be contributing to the increased transmissibility of the Delta variant. Therefore, it is important to understand both the RBD-ACE2 binding as well as the RBD-Ab binding.

In this study, we investigate the effects of the mutations in the Delta variant on the structure of the receptor-binding interface of the RBD as well as the RBD-ACE2 interactions and RBD-neutralizing Abs interactions. We examine the SARS-CoV-2 Ab-RBD complexes available in the protein data bank (PDB) and compare the differences in the RBD-Ab interactions due to the mutations in the Delta variant. Our results suggest that the Delta variant features a stable but slightly reorganized receptor-binding interface that can lead to weakened interactions with some neutralizing Abs resulting in immune evasion.

## 2. Materials and Methods

### 2.1 System Preparation

The Delta variant RBD was prepared by introducing the L452R/T478K mutations to the WT structure taken from the CHARMM-GUI^34, 35^ COVID19 repository (spike protein-ACE2 complex, PDB ID 6VSB, 6VW1)^36–38^. The RBD-only system for the Delta variant was set up and simulated the same way as for the WT, B.1.1.7, B.1.351 variants reported in our earlier work in Bhattarai et al.^39^. The RBD-ACE2 complex for the Delta variant was modeled by superimposing the Delta RBD (the last frame of the 600 ns molecular dynamics simulation) onto the WT RBD-ACE2 complex, with the receptor-binding interface of the RBD selected for the structural alignment.

The structures of the RBD-Ab complexes were retrieved from the RCSB Protein Data Bank (PDB)^37^. Four representative RBD-Ab systems were prepared for simulation: two for the WT (pdb IDs 6xe1 and 7chb) and two for the Delta variant. The RBD-Ab complexes for the Delta variant were modeled by superimposing the RBD from the last frame of the 600 ns Delta simulation onto the WT RBD-Ab complexes. All systems were solvated in TIP3 water molecules in 0.15 M salt concentration using CHARMM-GUI.

### 2.2 Molecular Dynamics Simulations

All-atom molecular dynamics (MD) simulations were performed with NAMD 2.14^40^ using the Charmm36m force field^41, 42^. The structures were equilibrated for 2 ns with a timestep of 2 fs after a short minimization. The production runs were performed under constant pressure of 1 atm, controlled by a Nose−Hoover Langevin piston^43^ with a piston period of 50 fs and a decay of 25 fs to control the pressure. The temperature was set to 303 K and controlled by Langevin temperature coupling with a damping coefficient of 1/ps. The Particle Mesh Ewald method (PME)^44^ was used for long-range electrostatic interactions with periodic boundary conditions and all covalent bonds with hydrogen atoms were constrained by ShakeH^45^. The Delta RBD-only system was run for 600 ns, whereas the RBD-ACE2 and RBD-Ab systems were run for 100 ns. For comparison with the WT, B.1.1.7 and B.1.351, 600 ns of simulation was performed for all three systems. Additional replicas of RBD Delta complex was run for 300 ns and the trajectories from our earlier work^39^ were analyzed for WT, B.1.1.7 and B.1.351 as additional replica runs. The simulations and systems in this work are summarized in Table S1. Visualization and analysis of the trajectories were done with Visual Molecular Dynamics (VMD)^46^.

### 2.3 Ab-RBD binding using MaSIF

To evaluate the Ab-binding to the RBD, we used a pre-trained *MaSIF-search* geometric deep learning model^47^. The model evaluates the protein-protein binding potential, given two surface regions (*patches*) from distinct proteins. Each patch is represented as the radial surface region of size 12Å, where each point in the mesh is associated with five geometric and chemical features: shape index, distance-dependent curvature, electrostatics, charge, and hydropathy. The deep learning model converts patches into 80-dimensional feature space, such that distance between embedded vectors from native binders is minimized. This distance is referred to as the *binding cost*, i.e., a lower output score represents a better interaction. The binding cost of the Ab-RBD complex was computed as the average of the output scores of three best patch pairs.

## 3. Results and Discussion

The RBD of the spike protein has mutations N501Y in B.1.1.7, K417N/E484K/N501Y in B.1.351, and L452R/T478K in Delta B.1.617.2. The RBD structure complexed with ACE2 is shown in Fig. 1 with the locations of the RBD mutations highlighted for the Delta variant. The ACE2 interacting interface of the RBD features an antiparallel β-sheet in the middle (residues 452-455, 492-495) and loop segments in the ends. The RBD mutations in or around the interfacial region can directly impact the RBD’s ability to bind ACE2 or neutralizing antibodies.

**Figure 1:**
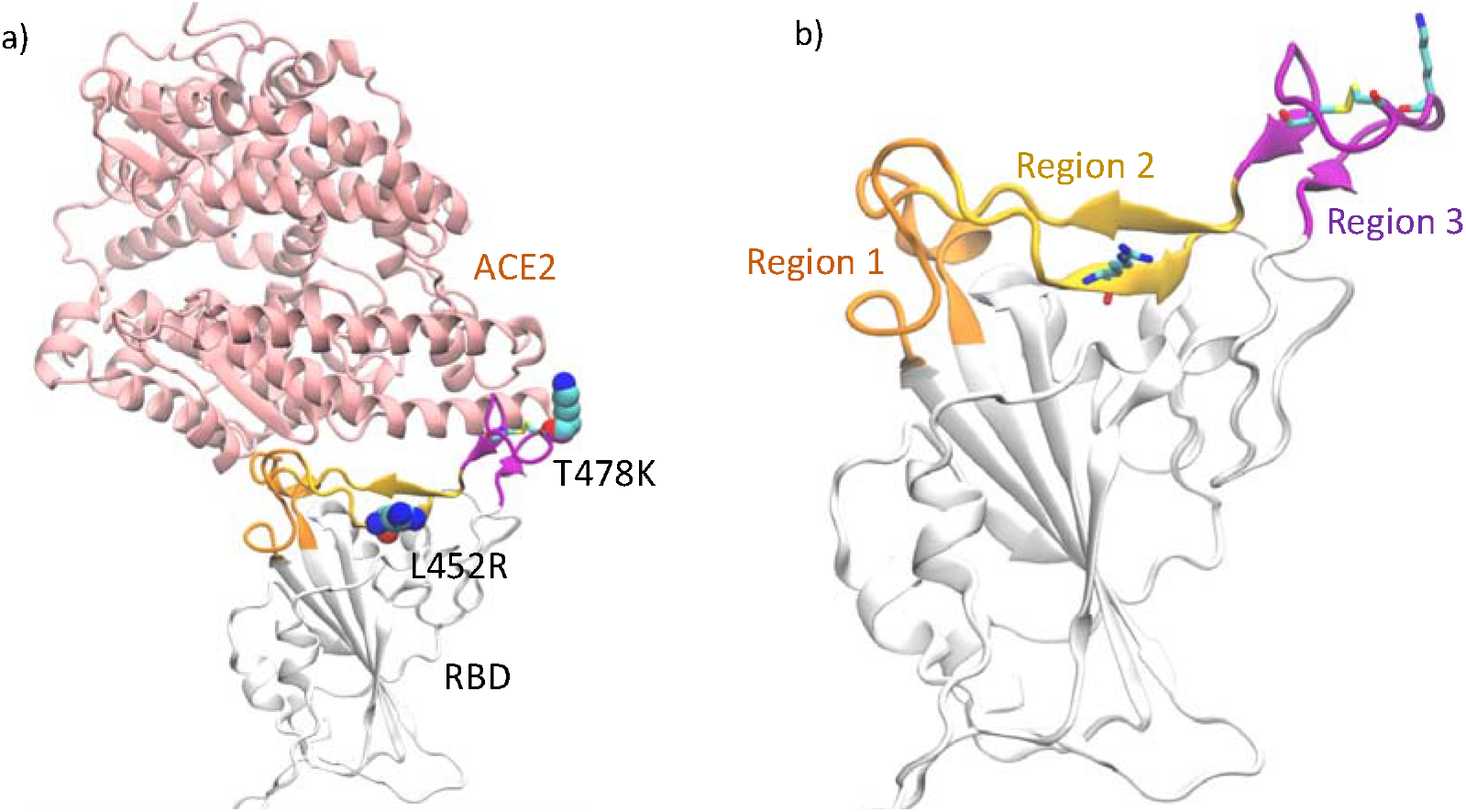
a) RBD complexed with ACE2. The locations of the mutations in the RBD of Delta variant are highlighted in VDW representation b) The loop segments consisting of residues 438-447 and 499-508 (Region 1) are highlighted in orange, the β-sheet region consisting of residues 448-455 and 491-498 (Region 2) are highlighted in yellow and the receptor-binding loop consisting of residues 472−490 (Region 3) in purple. The disulfide bond in the loop as well as the mutations in the Delta variant are shown as sticks.

### 3.1 Structural changes due to mutation in Delta variant

To investigate the RBD dynamics and the structural changes due to mutations, we performed MD simulations of the RBD of the Delta variant B.1.617.2 and compared the results with the WT, B.1.1.7, and B.1.351 variants. Both of the mutations, L452R and T478K, in the Delta RBD are in the receptor-binding interface comprised of a motif spanning residues 438 to 508. The same interface is a target for many neutralizing antibodies. Therefore, any changes in the receptor-binding interface can affect both the receptor binding to the host ACE2 as well as Ab-binding. To assess the structural changes in this interface, we analyzed different regions of the interface as shown in Figure 1. We calculated the root mean square fluctuations (RMSF) of the RBD and plotted the results in Fig. S1, which shows that the amino acid residues in the β-loop-β motif (Region 3, residues 472-490) have the largest flexibility for all variants. The loop segments in Region 1 are not found to change significantly compared to WT, and therefore we focused on the β-sheet in Region 2 and the β-loop-β motif of Region 3.

#### 3.1.1 Structural rearrangements in the interfacial beta sheet region

Figure 2a shows the β-sheet region of the receptor-binding motif (RBM) interface (Region 2 comprised of residues 448-455, 491-498) containing a hydrogen-bond network (Fig. 2b) that creates a stable interface. In the WT, residues in each segment have β-secondary structure (β5: 452-455 and β6: 491-495)^48^ with backbone hydrogen-bonds, whereas the additional residues in each segment (448-451 and 496-498) are mostly unstructured. Figure 2b shows the hydrogen-bonding in the RBM in the WT and the Delta B.1.617.2 variant at the end of 600 ns MD simulations. The hydrogen-bond analysis of Region 2 of the interface (Fig. 2c) shows that WT and B.1.1.7 have similar H-bond interaction patterns, whereas B.1.351 and B.1.617.2 have noticeably different H-bond patterns.

**Figure 2:**
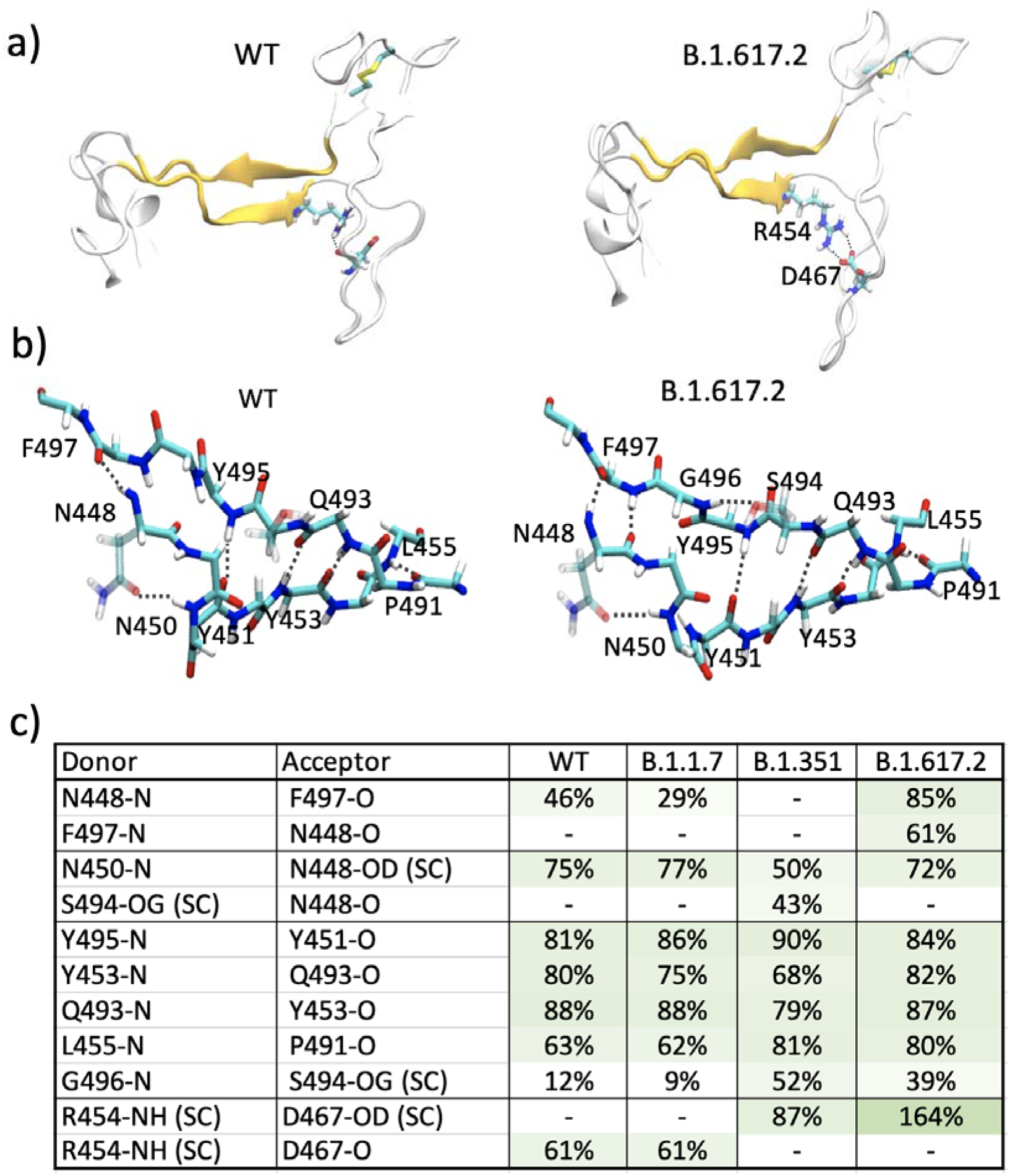
a) RBM showing the antiparallel β-strands. Residues R454 and D467 participating in ionic interactions in the Delta variant are shown as sticks. b) Hydrogen-bond network in the β-sheet region of the RBM for the WT and the Delta variant. c) % hydrogen bond occupancy obtained from the last 300 ns for the interactions in WT, B.1.1.7, B.1.351, and B.1.617.2. The sidechain interactions are denoted as SC.

Interestingly, the analysis of Fig. 2c shows that a slight reorientation of residue G496 in the Delta variant results in much stronger hydrogen bonding between the β-strands. Most notably for the Delta variant, 1) a more stable network of backbone hydrogen bonds is established, with a new hydrogen bond N448(N)-F497(O) formed, and 2) a significantly enhanced salt-bridge interaction between the R454 side chain and D467 side chain is observed. These two changes appear to be due to the mutation L452R which gives a slightly enhanced β-structural propensity^49, 50^ in β5. It has recently been shown that the L452R mutation in another variant of concern, B.1.427/B.1.429, caused reduction in nearly half of the tested monoclonal Abs.^32^, highlighting the dangerous consequences of this mutation. We analyzed 300 ns re-runs for each of these variants and compared in Fig. S2. Although the R454-D467 backbone hydrogen bond has not switched to a side chain interaction by 300 ns in the re-run of the Delta variant, the presence of the N448(N)-F497(O) hydrogen-bonding between the β-strands is consistent (Fig. S2). As seen in Figs. 2b and 2c, the interactions in the antiparallel β-strands are enhanced in Delta B.1.617.2, thereby stabilizing the receptor-binding interface. This may directly affect how the RBD binds with ACE2 and with neutralizing antibodies. We note that the recently solved crystal structure of the L452R variant B.1.617.1 (PDB ID 7orb) does not show these hydrogen bonds, suggesting that the changes observed here are perhaps a result of a dynamic reorganization. Similarly, almost all RBD structures show that R454 side chain makes hydrogen bonds with backbones of D467 and/or S469 but some structures do show possible side chain interactions with D467 (e.g. PDB ID 7n1q^33^, 7kdj^51^), suggesting an agile network of hydrogen-bonding in this region.

#### 3.1.2 Structural rearrangements in the β-loop-β motif

The flexible β-loop-β motif (Fig.1, Region 3, residues 472-490) contains a disulfide bond between resides C480-C488, and the mutation T478K in the Delta variant also lies in this loop. We explored the mutation-induced changes in the flexibility of this region, and rearrangements in the hydrogen bonding for the different variants (WT, B.1.1.7, B.1.351, B.1.617.2). We find that the Delta variant features a significantly different loop structure. While all variants have a flexible loop in this region^39^, the Delta variant shows a reduced flexibility (Fig. S1) as it adopts a more stable yet different conformation compared to other variants. The difference in the conformational change in the loop can be seen from the changes in the disulfide bond dihedral angle C-C_α_-C_β_-SG for C480 as displayed in Fig. 3a. Compared to the WT, the dihedral angle as a function of time for the Delta variant shows a quick flip early and then remains stable in a new orientation.

**Figure 3.**
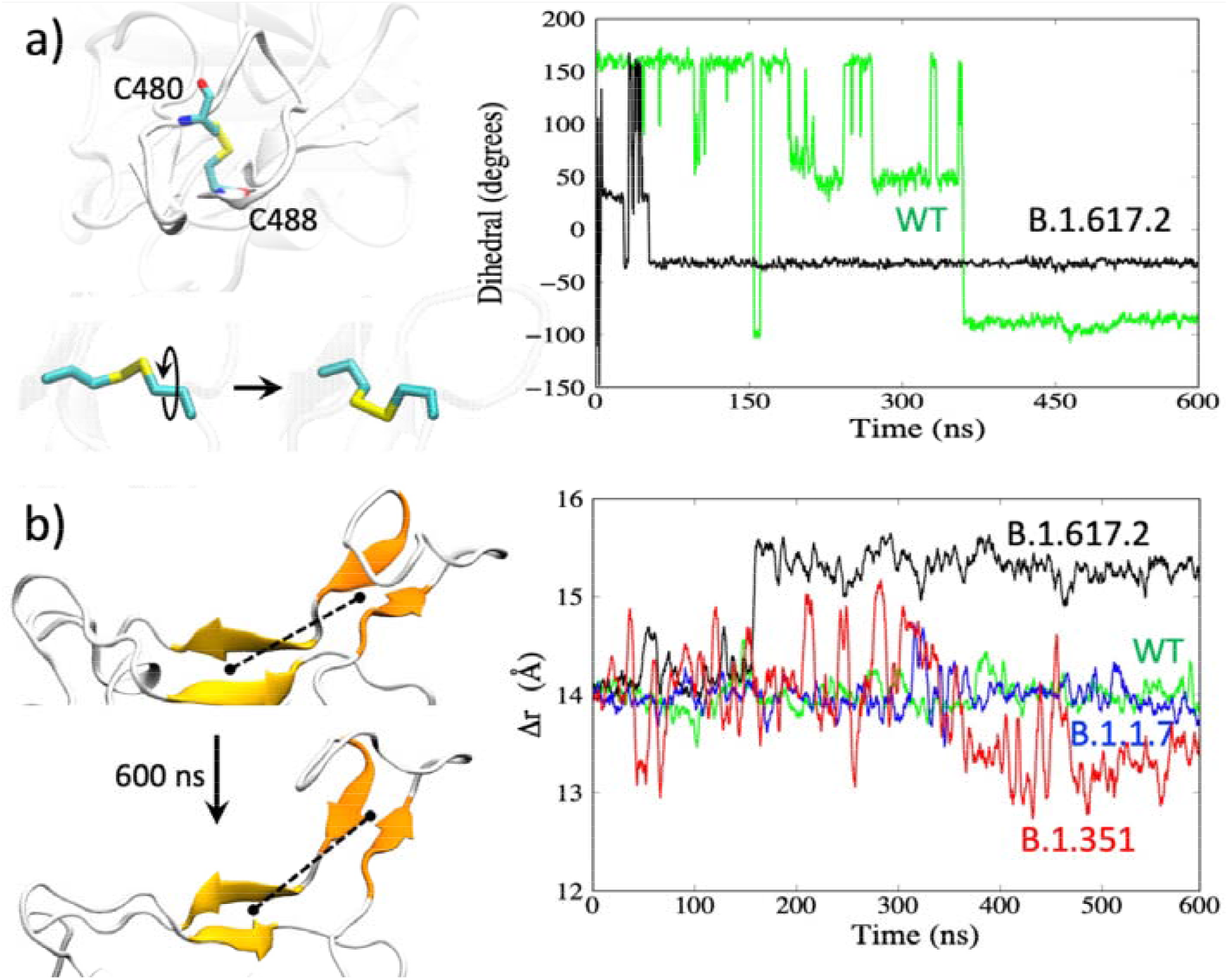
a) Reorientation of the disulfide bond. Right: changes in the dihedral angles for WT and the Delta variant. b) Distance between the center-of-mass between the two β-sheets (shown as the dotted line on the left).

The combination of the changes in the β-sheet region (Fig. 1b, Region 2) and the β-loop-β motif (Fig. 1b, Region 3) appears to result in an overall change in the receptor-binding interface. In the Delta variant, Regions 2 and 3 are farther apart. This is shown by the separation distance (dotted lines in Fig. 3b) between the center-of-mass (COM) of the two β-strands in Region 2 (Fig. 1b: β-strands 452-455 and 492-495) versus the COM of the two β-strands in Region 3 (Fig. 1b: 472-475 and 487-490). The distance plot in Fig. 3b (right) shows a stable but slightly extended Region 2 – Region 3 receptor-binding interface for the Delta variant (black curve) with a 1.5 Å increase in the COM separation distance. Overall, these structural changes and differences in loop flexibility can impact the ACE2 and Ab-binding. The reduced fluctuations in the Delta RBD with an altered receptor-binding interface could result in weaker interactions with neutralizing antibodies leading to immune evasion.

### 3.2 Antibody binding to the Delta RBM and possible mechanism of immune evasion

A recent study illustrated the mechanism of immune evasion by a variant of concern B.1.427/B.1.429^32^. Specifically, the same mutation found in the Delta variant, L452R, was responsible for reduced neutralizing activities in many of the monoclonal Abs tested, whereas re-grouping of a disulfide bond in a different RBD site caused the loss of activities for all Abs tested. To assess the impact on the Ab binding due to the changes in the receptor-binding interface caused by the mutations in the Delta RBD, we first examined the interfacial interactions in the Ab-RBD complexes available in the protein data bank (PDB) in the WT. Of the 118 RBD-Ab complexes with Ab bound in the receptor-binding interface retrieved from the Protein Data Bank, 47 non-repeating complexes were considered for further analysis. The Ab-RBD complexes (pdb IDs) are listed in Table S2 and visualized in Movie S2. We identified the RBD residues involved in ionic or hydrogen bond interactions in each complex and plotted in Fig. 4 the frequency of occurrences of the important residues in all complexes. While this distribution may be inherently biased due to the available pdb structures of the complexes, it provides a general idea of the preferred interfacial RBD binding epitope sites for a sample of Abs. From Fig. 4, we see that the majority of the Abs have interactions with the β-loop-β residues Y473, A475, N487, E484, among others.

**Figure 4.**
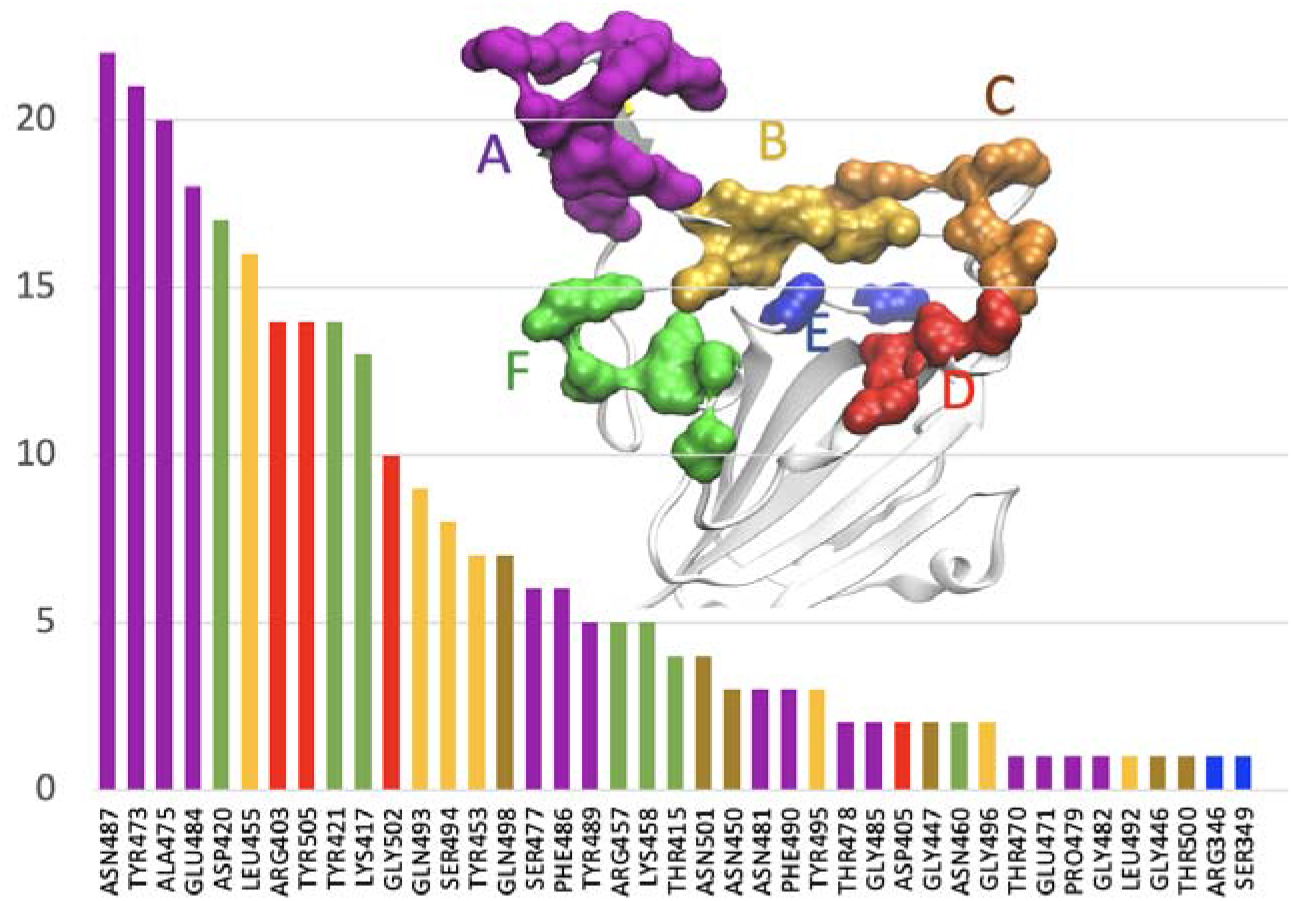
Frequency of occurrences of the RBD residues involved in hydrogen-bonding with Ab in 47 complexes from the Protein Data Bank.

Inspection of the Ab-RBD complexes for the WT shows that many of the Abs anchor at multiple sites. For example, in many Ab-RBD complexes, including 6xe1, 7b3o, 7cdi, and 7cjf Abs bind at A475/G485 at one site (site A in Fig. 4) and R457/K458 (site F in Fig. 4) at another site, as shown in the figure. We grouped different sites and color-coded as shown in Fig. 4. To examine how the changes in these sites may affect the Ab binding, we plotted the C_α_ distance between the residues K458 and A475 belonging to two Ab-binding sites in Fig. 5 for both the WT and the Delta variant. As shown in Fig. 5b, the distance between these residues (458-475) mostly remains at ∼9 Å for the WT. However, the same (458-475) distance for the Delta variant in Fig. 5c increases to ∼14 Å by 150 ns and remains stable at that distance. This 4-5 Å increase in the Ab-binding sites suggests that the Ab-binding will be severely affected, and the Ab becomes insensitive to Delta RBD binding. While ACE2 binds at the site of 475/487 (site A in Fig. 4), it does not bind at the site of 457/458 (site F) and therefore the increase in the K458-A475 distance does not affect the ACE2 binding. Instead, ACE2 binds at 475/487 and Q493 (in the middle of β6). Therefore, we also plotted the distance between the residues N487 and Q493. Interestingly, despite the structural changes, this distance in both the WT and the Delta variant remains nearly the same (16-17 Å) as seen in Fig 5b and 5c. This suggests that the structural changes have not affected the ACE2 binding sites but significantly affected the Ab-binding sites, suggesting a possible immune evasion by the Ab while maintaining the ability of receptor binding.

**Figure 5.**
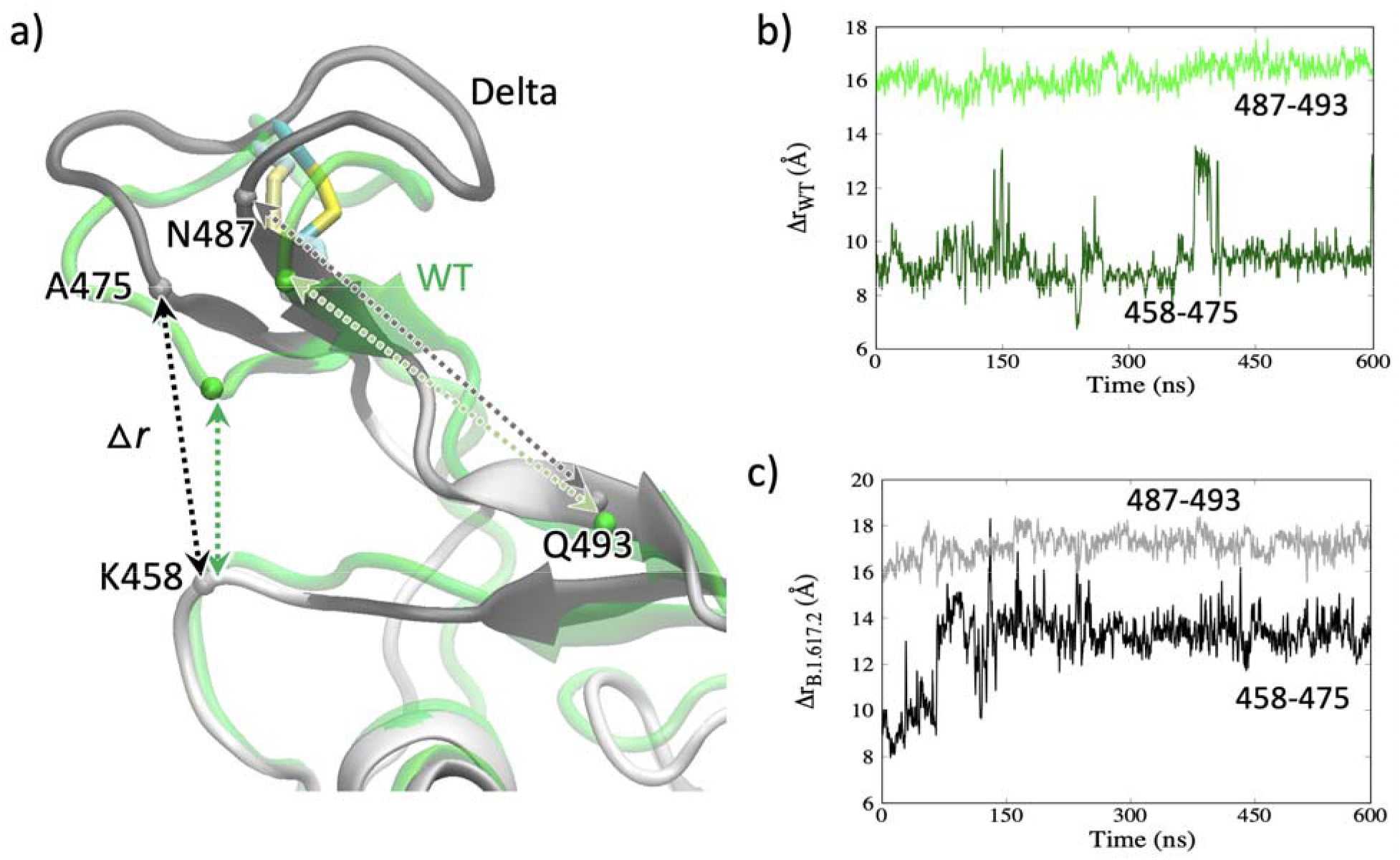
a) Amino acid residues involved in Ab-binding or ACE2-binding. b) The C_α_-C_α_ distances between the residue pairs K458-A475 and N487-Q493 in WT. c) The C_α_-C_α_ distances in the Delta variant.

### 3.3 ACE2 binding vs. antibody binding in the Delta variant

With the observation of the increase in the distance between the two sites in the Delta RBM, we investigated how the changes affect ACE2 and Ab binding. If the ACE2 binding is maintained or enhanced but the Ab binding is weakened, at least for a set of neutralizing Abs, that would mean that the virus is less sensitive to the Abs thereby making it more effective at infecting and spreading. To explore this, we performed simulations of the RBD-ACE2 complex and Ab-RBD complexes for the Delta variant and compared with the complexes of the WT. Since the complexes for the Delta variant were modeled from the RBD obtained from the 600 ns simulation, the interactions are expected to evolve, whereas those in the WT remain steady. The hydrogen bond interactions in the RBD-ACE2 as well as the Ab-RBD complexes are shown in Fig. 6. In the WT RBD-ACE2 complex, the RBD residues that primarily participate in hydrogen-bond interactions include K417, Y489, G502, E484, T500 and N487. The RBD residue K417 forms a strong salt-bridge with D30 of ACE2 in WT and B.1.1.7 but not in B.1.351 due to the K417 mutation^39^. These WT interactions are still present in the complex with the Delta RBD, though the % occupancy are reduced (Table S3). With some of the major interactions, including the K417-D30 salt-bridge, still present in the complex, ACE2 binding seems tolerate the structural changes in the Delta RBD.

**Figure 6.**
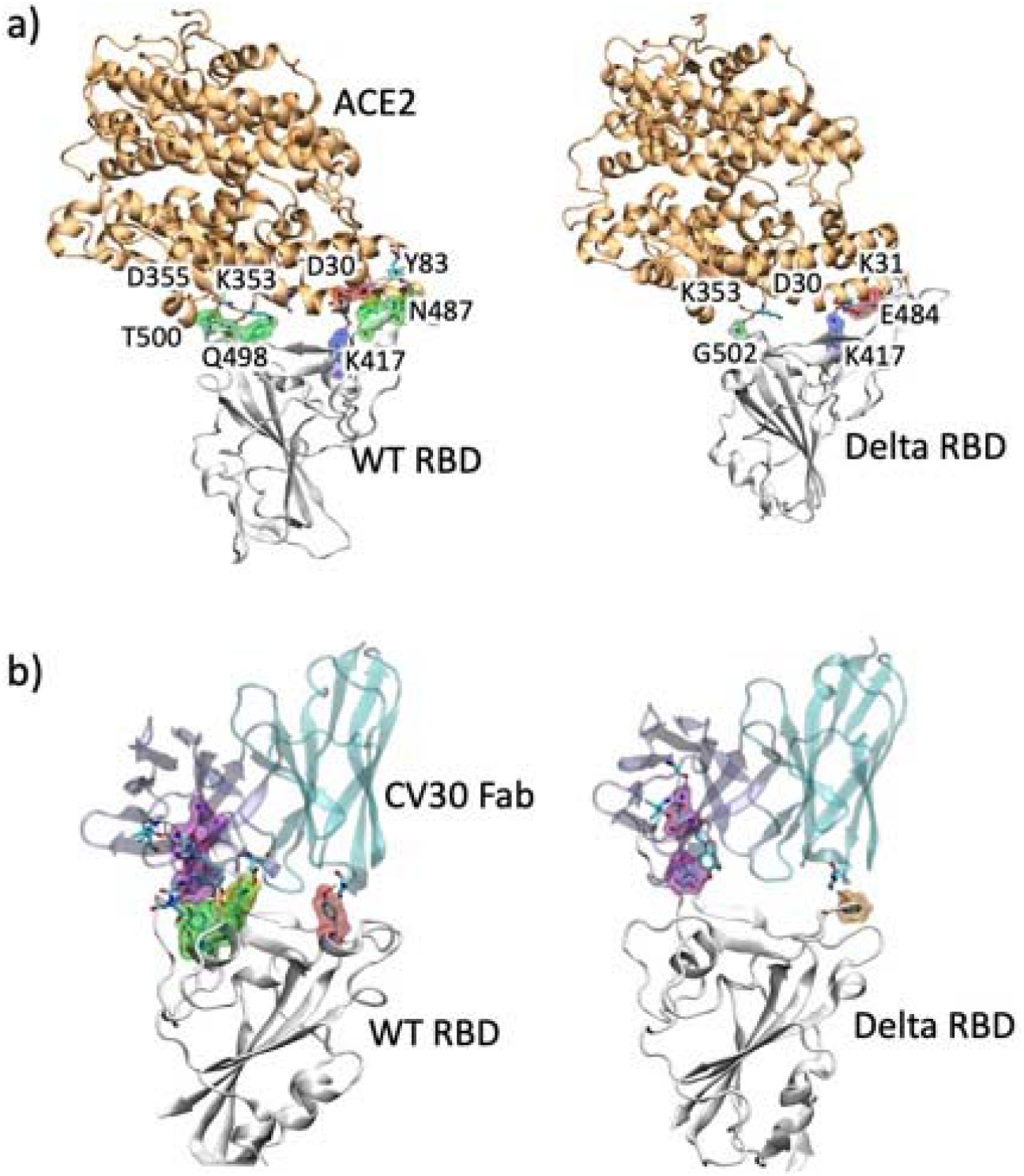
a) Structures of the ACE2-RBD complexes for WT and the Delta variant at the end of the 100 ns simulations. b) Ab-RBD complex for WT and the Delta variant, with CV30-Fab neutralizing Ab (PDB 6xe1) complexed with RBD. The interacting sites are highlighted in surface representation for the RBD and sticks for the ACE2 or Ab.

We next compared the Ab binding in the WT and the Delta RBD. We considered two examples of the Ab-Delta RBD complexes modeled from the WT RBD complexed with the neutralizing Ab CV30 Fab (pdb ID 6xe1)^52^ and complexed with the Ab BD-236 Fab (pdb ID 7chb)^53^. The 100 ns simulation of the CV30-Delta RBD model shows a less stable complex with significantly reduced interactions. As shown in Fig. 6b, Ab in the WT has interactions with residues in three clusters that are intact during the simulation. However, in the Delta RBD, the Ab is only able to bind at site A or F but not both. This is consistent with the argument made on the basis of Fig. 5. The major hydrogen bonds, including those with Y473, Y421, L455, A475, R457, R403, K417, and Y505 in WT are broken or weakened in the Delta variant. The Ab-RBD complex modeled from 7chb (complex with BD-236 Fab) also shows reduction in the number and strength of the hydrogen bonding. Some of the major hydrogen bonds withY453, G502, D420, and K417 in the WT BD-236 complex are still present in the Delta variant during the 100 ns simulation (Table S4). Figure 6b displays the final conformations of the Ab-RBD complexes for the CV30 Ab, with the RBD residues colored according to the grouping in Fig. 4. RBD structures complexed with BD-236 for the WT and the Delta variant are given in Fig. S3, and the residues and the % hydrogen bonding occupancies are given in Table S4. Binding analyses using *MaSIF-search*^47^ shows that both Ab-RBD complexes in the Delta variant have significantly reduced binding (Fig. S4). The violin plot in Fig. S4 show the distribution of the binding costs extracted from the Ab-Protein complexes in the SAbDab structural antibody database^54^. The scores for the WT complexes lie within the distribution but the complexes for the Delta show significantly higher binding costs. While Abs that bind at other sites may not be affected, Abs that bind at A-F can become insensitive, albeit differentially, to the Delta RBD binding.

## 4. Conclusion

The Delta variant (B.1.617.2) of the SARS-CoV-2 has become one of the most worrisome variants so far during the pandemic and is rapidly spreading worldwide, making it responsible for the recent surges in infections and deaths. While current vaccines are still shown to be protective against this variant, it is also becoming clearer that it can escape the immune system by making neutralizing Abs from prior infections or elicited by vaccines less sensitive to binding with the spike protein. In this work, we performed molecular dynamics simulations of the Delta variant RBD with the mutations L452R/T478K and investigated the resulting structural changes in the receptor- and Ab-binding interfaces. We find that the Delta variant presents a noticeably different receptor-binding interface compared to the WT, B.1.1.7, and B.1.351. Specifically, the receptor-binding β-loop-β motif adopts an altered conformation which appears to cause shifts in the Ab-binding epitope regions that can reduce the binding affinities for some neutralizing Abs. We investigated this by performing simulations of two Ab-RBD complexes and found that one of the complexes shows significantly reduced interactions between the Ab and the RBD, suggesting a possible mechanism of the immune escape by the Delta variant. Even though the Ab-resistant conformations obtained in these simulations may represent only a subset of the conformational ensemble, they can still contribute considerably to the reduced sensitivity of the Abs. Future work with a full mapping of the conformational space of the receptor-binding interface may shed further light on the nature of the interactions with the common anti-RBD Abs, providing useful information on vaccine efficacies. Understanding how the structural changes alter the RBD’s ability to present itself in the up conformation in the spike trimer or its ACE2 binding affinity can also inform us on the variant’s transmissibility.

## Supporting Information

Additional figures showing the hydrogen-bonding residues in the complexes and tables listing the Ab-RBD complexes and the interacting residues.

Movie S1: showing the 600 ns trajectory of the Delta variant RBD.

Movie S2: showing a sample of Ab-RBD complexes from the protein data bank.

## Acknowledgements

This work was supported by the National Science Foundation under Grant No. 2037374. NB acknowledges the Dissertation Year Fellowship support from the University Graduate School at Florida International University. The authors thank the COVID-Informatics research team at Florida International University for helpful discussions.

