## Supplemental File for "Mutation-induced Changes in the Receptor-binding Interface of the SARS-CoV-2 Delta Variant B.1.617.2 and Implications for Immune Evasion"

**Supporting Information**

Table S1: Details of the simulation systems.

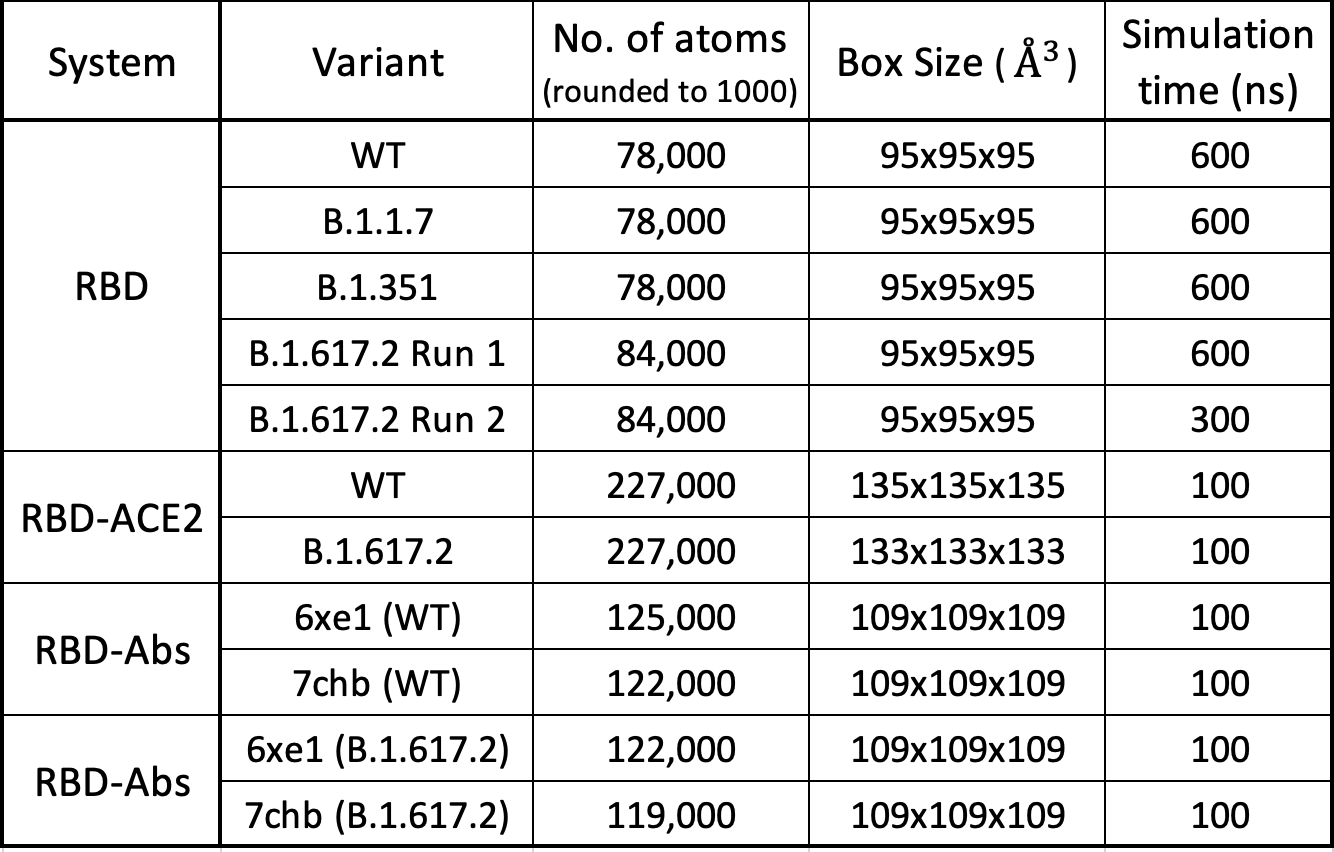

Note: Additional 300-ns re-runs (not listed here) for the WT, B.1.1.7, and B.1.351 were

analyzed from our earlier work (Bhattarai et al. J. Phys. Chem. B. 125,7101, 2021).

Table S2: List of the Ab-RBD complexes obtained from the Protein Data Bank. The representative structures from each of the non-repeating groups considered for structural analysis are highlighted in boldface. The structures from the same family of complexes are underlined.

**6xc2**, 6xc3, 6xc4, 6xc7, **6xdg**, **6xe1**, **6xey**, **6xkp**, **7b3o**, **7beh**, 7bei, 7bej, 7bek, 7bel, 7bem, 7ben, 7beo, 7bep, **7bwj**, **7bz5**, **7c01**, **7cdi**, 7cdj, **7ch4**, 7ch5, **7chb**, **7chc**, 7che, 7chf, 7chh, **7cho**, 7chp, 7chs, **7cjf**, **7cm4**, 7cwl, 7cwm, 7cwn, **7cwo**, 7cwu, **7czp**, 7czq, 7czr, 7czs, 7czt, 7czu, 7czv, 7czw, 7czx, 7czy, 7czz, 7d00, **7d03**, 7deo, **7deu**, **7dk4**, 7dk5, 7dk6, 7dk7, **7e23**, **7jmo**, 7jmp, **7k43**, 7k45, 7k4n, **7k8m**, 7k8s, 7k8t, 7k8x, **7k90**, **7k9z**, **7kfv**, 7kfw, 7kfx, 7kfy, **7klg**, 7klh, 7kmg, 7kmh, **7kmi**, 7kmk, **7kml**, **7kn6**, 7kn7, **7ks9**, **7kxj**, 7kxk, **7kzb**, **7l3n**, **7l56**, 7l57, 7l58, 7l5b, **7laa**, **7ljr**, **7lop**, **7m6d**, 7m6f, 7m6h, 7m6i, **7mf1**, **7mjj**, 7mjk, 7mjl, **7nd4**, 7nd5, 7nd6, 7nd7, 7nd8, 7nd9, 7nda, 7ndb, **7neg**, 7neh, **7nx6**, 7nx7, 7nx8, 7nx9.

Table S3: Hydrogen bond details analysis of RBD-ACE2 for WT and B.1.617.2.

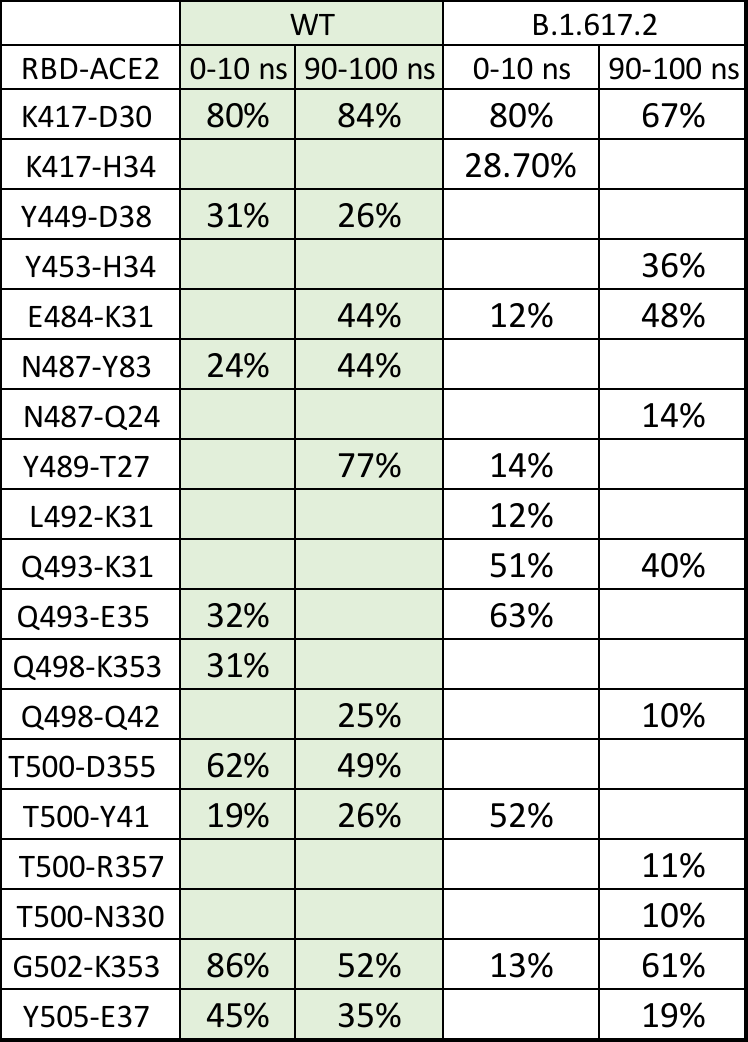

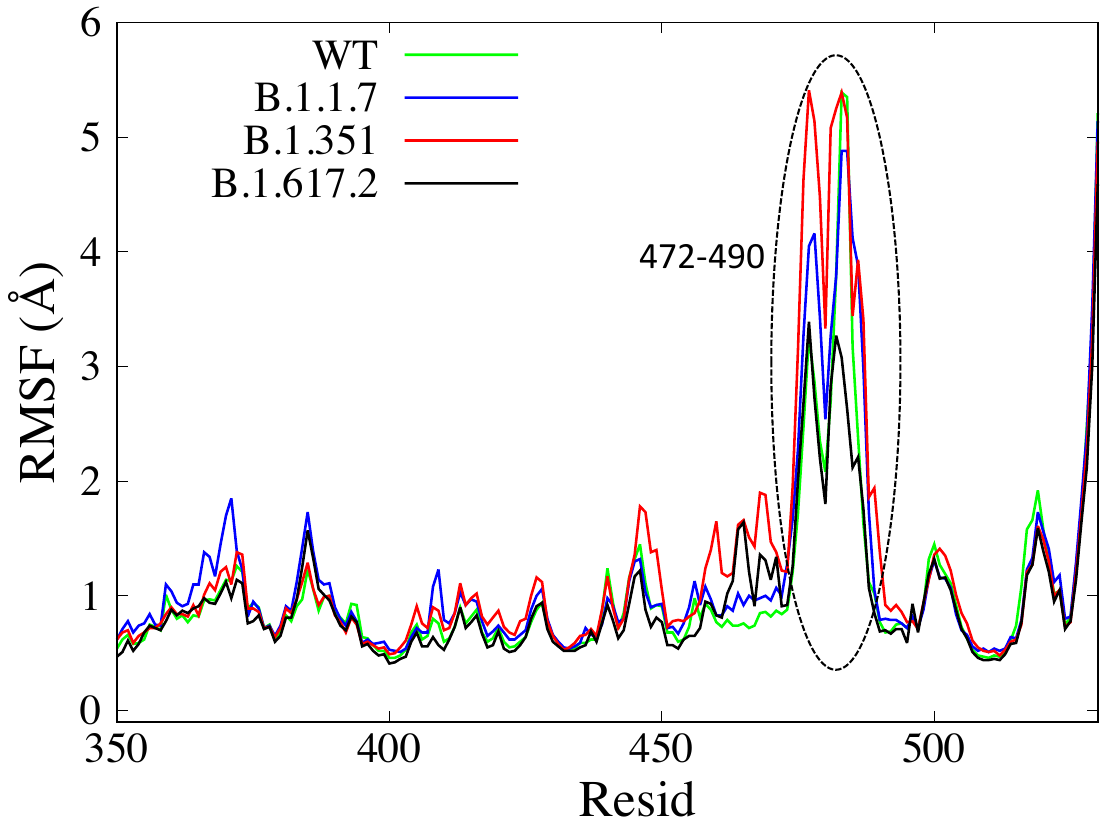

Figure S1: Root Mean Square Fluctuations (RMSF) of amino acids calculated from the last 300 ns of MD simulations. The flexible β-loop-β region at the RBM interface is indicated with a dotted ellipse.

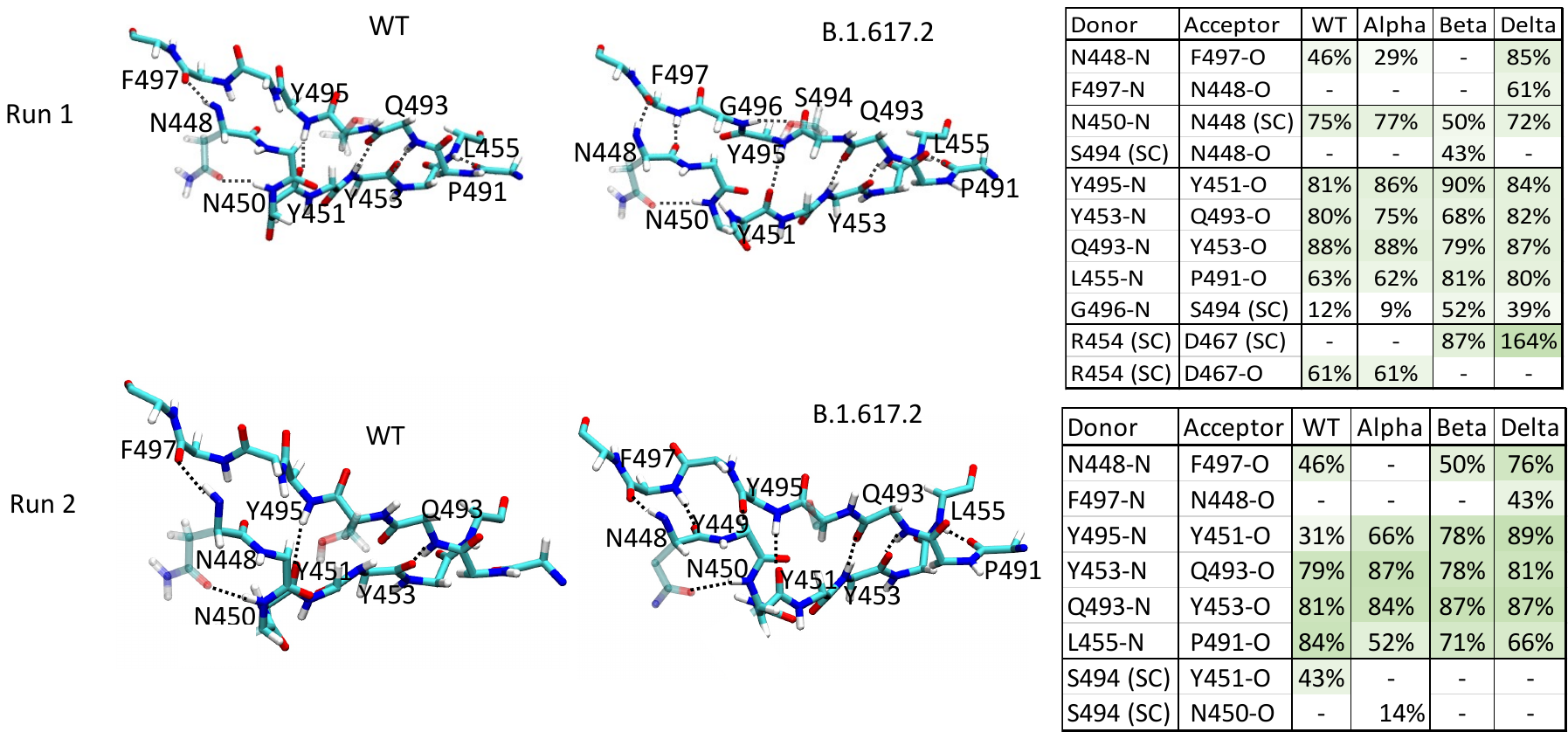

Figure S2: Hydrogen bonding network in beta sheet at the interface for 600 ns MD simulations (run 1) and rerun for 300 ns (run2). The backbone hydrogen bonding between the β-strands for the Delta variant is consistent in both run 1 and run 2.

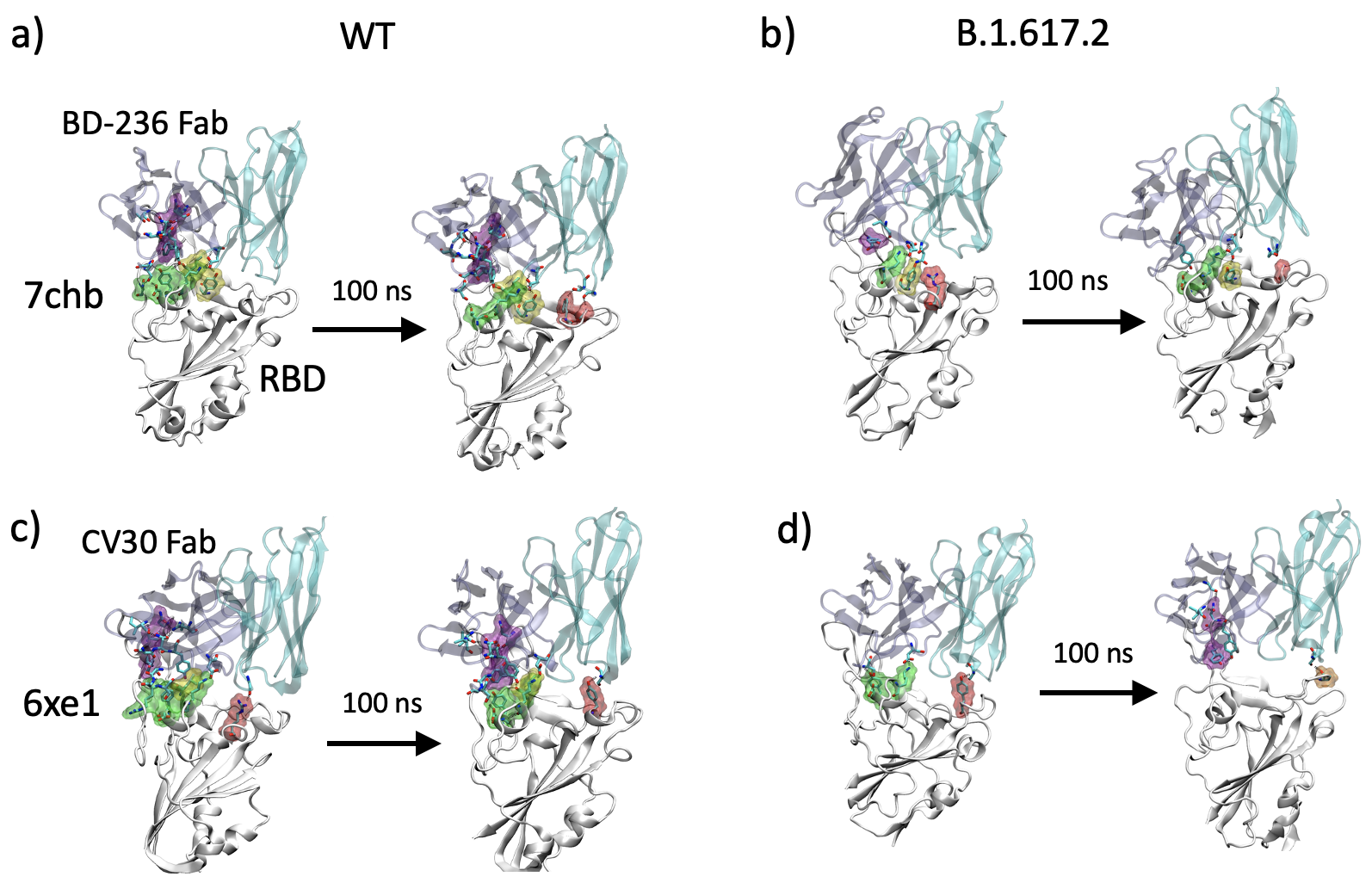

Figure S3: Structures of the ACE2-RBD complexes for WT and the Delta variant at the end of the 100 ns simulations. a) WT BD236-RBD complex (pdb ID 7chb), b) Delta BD236-RBD complex c) WT RBD-CV30 complex (PDB 6xe1), and d) Delta RBD-CV30 complex. The interacting sites are highlighted in surface representation for the RBD and sticks for the Ab.

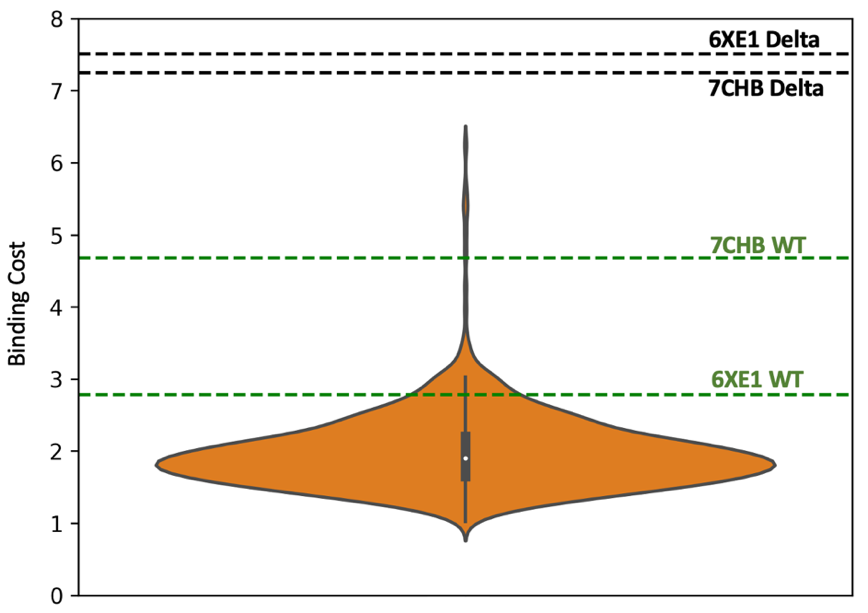

Figure S4: Violin plot showing the distribution of the binding cost of Ab-Protein complexes from the SAbDab database. Lines represent the binding cost values for WT (green) and Delta (black).
Table S4: Hydrogen bond details for the RBD complexed with a) CV30 Fab antibody (PDB 6xe1) and b) BD-236 Fab antibody (PDB 7chb) for WT and B.1.617.2.

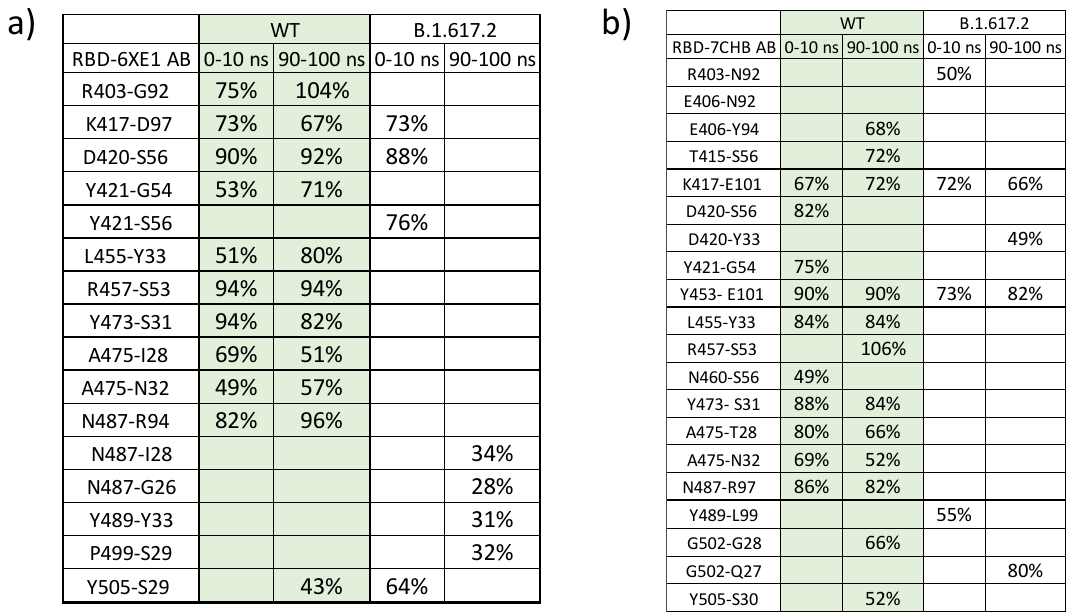

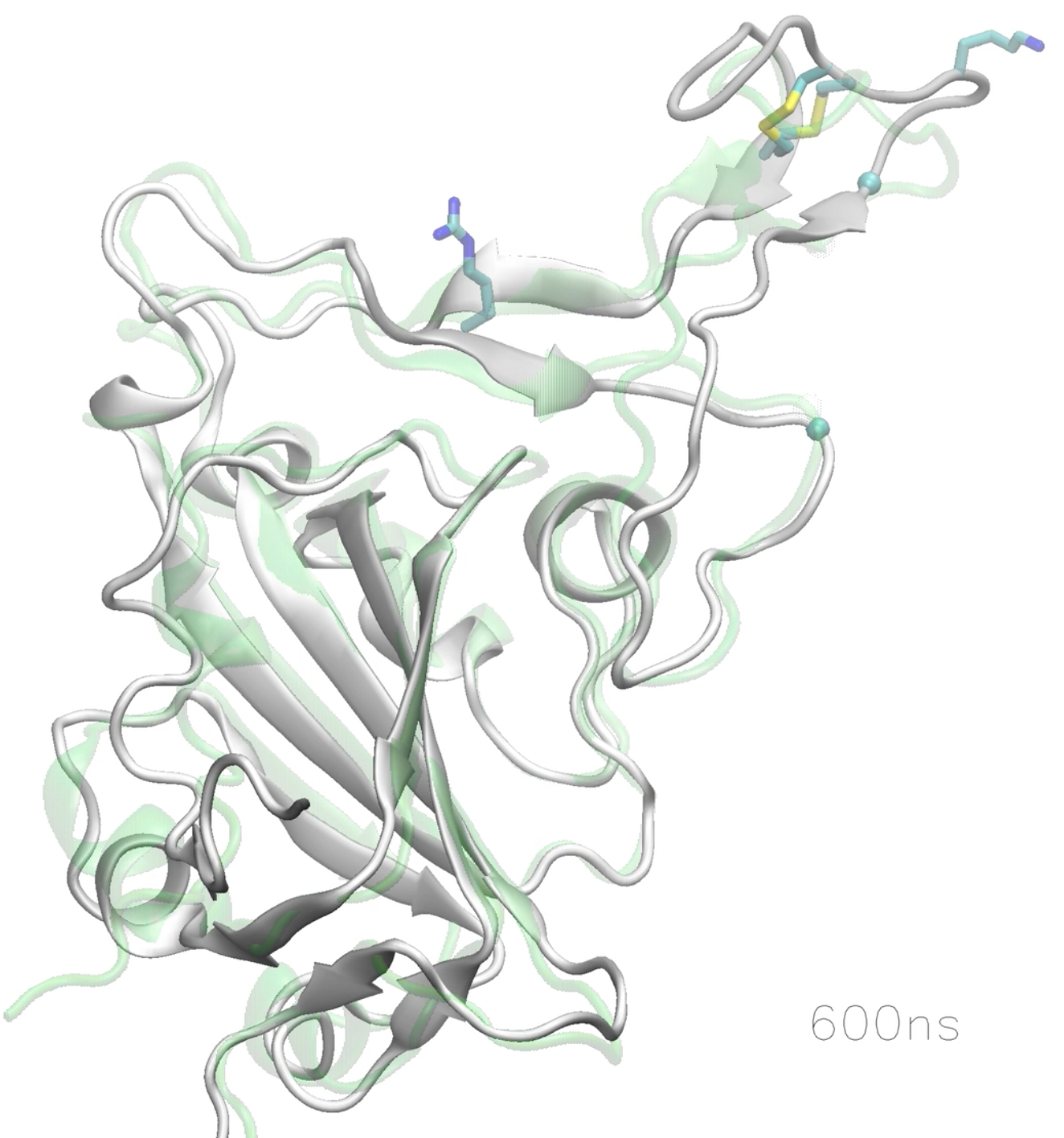

Movie S1: Dynamics of the Delta RBD during the 600 ns of MD simulation. The initial structure is colored light green (which is also the same structure for the WT) while the trajectory of the Delta RBD is shown in gray. The mutations L452R/T478K) as well as the disulfide bond in the RBM loop are shown as sticks.

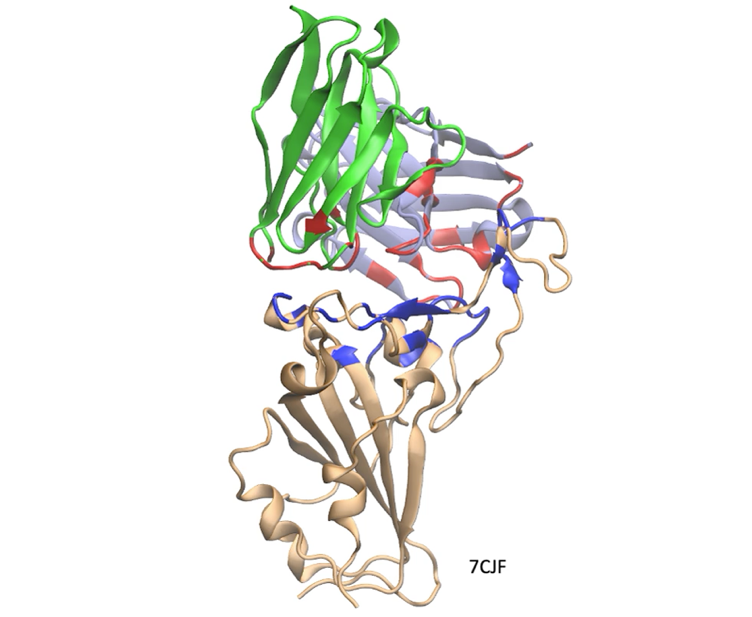

Movie S2: Various RBD-Ab complexes from the Protein Data Bank showing the interacting regions. The RBD is colored orange, the antibody chains as green and ice blue. The Ab residues within 3.5 Å of the RBD are highlighted in red and the RBD residues within 3.5 Å of the Ab are highlighted blue. The PDB Id for each complex is also displayed.
